# Life-long telomere attrition predicts health and lifespan in a large mammal

**DOI:** 10.1101/2020.06.22.165563

**Authors:** Luise A. Seeker, Sarah L. Underwood, Rachael V. Wilbourn, Jennifer Fairlie, Hannah Froy, Rebecca Holland, Joanna J. Ilska, Androniki Psifidi, Ainsley Bagnall, Bruce Whitelaw, Mike Coffey, Georgios Banos, Daniel H. Nussey

## Abstract

Telomere length measured in blood cells is predictive of subsequent adult health and survival across a range of vertebrate species. However, we currently do not know whether such associations result from among-individual differences in telomere length determined genetically or by environmental factors early in life, or from differences in the rate of telomere attrition over the course of life. Here, we measured relative leukocyte telomere length (RLTL) multiple times across the entire lifespan of dairy cattle in a research population that is closely monitored for health and milk production and where individuals are only culled in response to health issues and less due to poor milk production than on purely commercial farms. Our results clearly show that the average amount of telomere attrition over an individual’s life, not their average or early life telomere length predicted when an individual was culled. Within-individual telomere length attrition could reflect environmental or physiological insults which may accumulate to predict individual health-span. We also show that animals with more telomere attrition in their first year of life were culled at a younger age, indicating that early life stressors may have a prolonged effect on adult life.

## Introduction

Telomeres are repetitive DNA sequences that cap the ends of eukaryote linear chromosomes (Blackburn & Gall, 1978; De Lange, 2005). They shorten with the number of cell divisions *in vitro* as well as in response to oxidative stress and critically short telomeres trigger a DNA damage response that leads to replicative senescence or apoptosis (Harley, Futcher, & Greider, 1990; Olovnikov, 1973; Watson, 1972). In the last decade or so, measures of average telomere length (TL) taken from blood samples have emerged as an exciting biomarker of health across disciplines including biomedicine, epidemiology, ecology and evolutionary biology (Aviv & Shay, 2018; Harrington & Pucci, 2018; Wilbourn et al., 2018). Considerable among- and within-individual variation in TL has been observed, with a general pattern of rapid telomere attrition during early life and a plateau or slower decline thereafter (Aubert & Lansdorp, 2008; Baerlocher, Rice, Vulto, & Lansdorp, 2007). Both genetic and environmental factors, particularly those associated with physiological stress, predict TL in humans and other vertebrates (Angelier, Costantini, Blévin, & Chastel, 2017; Muhammad Asghar et al., 2015; Cherkas et al., 2006; Dugdale, Richardson, & Richardson, 2018; Epel et al., 2004). TL has also been repeatedly associated with health outcomes and subsequent survival in a variety of species, particularly humans and birds (Boonekamp, Simons, Hemerik, & Verhulst, 2013; Wilbourn et al., 2018) and also experimentally elongated TL in mice were associated with a survival advantage (Muñoz-lorente, Cano-martin, & Blasco, 2019). However, a major outstanding question remains to what degree associations between TL and health arise from constitutive differences in TL among individuals set by genes or early life conditions, or from the pattern of within-individual change in TL across individuals’ lives.

Estimates of the individual consistency of TL over time in both human and avian literature vary considerably among studies. Some studies report very high correlations within individuals across follow-up measurements (r > 0.8; (Benetos et al., 2013; Boonekamp, Mulder, Salomons, Dijkstra, & Verhulst, 2014)) and high heritability of TL (>0.7; (Atema et al., 2015; Broer et al., 2013)), and have shown in humans that the rank order in TL among individuals remains relatively consistent during adult life (Benetos et al., 2013). This implies most of the variation in blood cell TL occurs at the among-individual level and is predominantly determined by genetics and early-life environment (Atema et al., 2015; Benetos et al., 2013). In stark contrast to this, a growing body of literature studying both humans and non-human vertebrates reports much lower individual repeatability and heritability (Dugdale & Richardson, 2018; Voillemot et al., 2012). These studies demonstrate very high levels of within-individual variation in TL and show that changes in TL across consecutive measurements are highly dynamic (Fairlie et al., 2015; Farzaneh-Far et al., 2010). Although on average TL attrition over time tends to be the norm, a growing number of studies have found that a substantial proportion of individuals show telomere elongation over time and it is now clear this cannot be ascribed solely to measurement error (Bateson & Nettle, 2016; Spurgin et al., 2018). A recent cross-sectional study has shown that amongst different avian and mammal species, those with a fast telomere attrition rate have a shorter life span but without investigating if telomere attrition rate within individuals is predictive of their life span within the population as well (Whittemore, Vera, Martínez-nevado, Sanpera, & Blasco, 2019). To date, few studies have directly compared the power of an individual’s average or early life TL versus the pattern of within-individual change in TL to explain variation in measured associated with health and fitness. In Seychelle warblers and Alpine swifts both telomere length and telomere attrition rate were associated with survival (Barrett, Burke, Hammers, Komdeur, & Richardson, 2013; Bize, Criscuolo, Metcalfe, Nasir, & Monaghan, 2009) while in jackdaw nestlings telomere length was not predictive of lifespan but early life telomere attrition-accelerated by early life stress-was (Boonekamp, Mulder, Salomons, Dijkstra, & Verhulst, 2014). Contrary, in a study on elderly humans, there was an association between telomere length and longevity, but not between telomere attrition and life span (Yuan, Kronström, Hellenius, Cederholm, & Xu, 2018). To our knowledge no studies so far has investigated the relationship between telomere attrition and life span with data spanning the entire typical lifespan of the studied individuals. Here, we use an exceptionally detailed longitudinal study of dairy cattle to show that a life-long tendency for greater telomere attrition, rather than average TL, predicts productive lifespan, a metric closely linked to long-term health.

We used samples and data collected as part of the long-term study of Holstein Friesian dairy cattle kept at the Crichton Royal Research Farm in Dumfries, Scotland (Veerkamp, Simm, & Oldham, 1994). This herd, consisting of around 200 milking cows plus their calves and replacement heifers, has been regularly monitored since 1973 for a broad range of measurements, such as body weight, feed intake, signs of disease (health events), milk yield, productive lifespan and reasons for culling (Veerkamp et al., 1994). The cows are separated into two distinct genetic groups (a group that has been strongly selected for high milk fat and protein yield, and a control group) and are randomly allocated to two different diets (a high forage, low energy diet vs. a low forage, high energy diet group) (Veerkamp et al., 1994). Blood samples are routinely collected from members of the herd, initially within 15 days after birth and then approximately annually thereafter. Productive lifespan, which is the age of an individual at culling, is recorded for every animal together with a reason for culling. In this and other commercial livestock herds, culling is a response to diseases such as reproductive problems, lameness and mastitis. Although productive lifespan in livestock is clearly not a comparable measure to longevity in study of humans or wild vertebrates, we argue that because age at culling is tightly linked to health, it can be considered as a proxy for health-span, over and above its obvious importance as a metric of agricultural productivity. We have previously reported for this herd that TL is moderately repeatable and heritable, but that individual slopes relating TL to age vary considerably among individuals (Seeker et al., 2018; Seeker et al., 2018). We have also reported that TL at the age of one year predicts productive lifespan in this population of dairy cattle (Seeker et al., 2018). Here, we test the relationships between lifetime average TL as well as lifetime change in TL with productive lifespan.

## Materials and Methods

### Animal population and data collection

At the Crichton Royal Farm, 200 milking cows plus their calves and replacement heifers are kept at any time. One half of the milking cows belong to a genetic line that has been selected for high milk protein and fat yield (S), while the other half is deliberately maintained on a UK average productivity level (C). Calves and heifers of both genetic lines are kept together. After first calving all cows are randomly allocated to a high forage (HF) or low forage (LF) diet. The LF diet is energy richer than the HF diet and whilst the LF cows are housed continuously, the HF cows graze over the summer months. All cows are milked three times daily and milk yield is recorded. In the present study, these measurements were used to calculate an average milk production in kg per cow including all started lactations and it is referred to this below as “average lifetime milk production” (Figure S1). Every day cows leave the milking parlour over a pressure plate which detects signs of lameness. Behaviour and health events are documented after visual detection by farm workers (Figure S2). At the end of the animal’s life its productive lifespan and a reason for culling are recorded. Productive lifespan is the time from birth to culling in days and is a proxy for the health span of the animal, because all animals that remain healthy enough to generate profit for the farmer remain in the herd. The most frequent reasons for culling were reproductive problems, mastitis, lameness and injuries caused by accidents (Figure S3). Routine blood sampling takes place initially shortly after birth and then annually in spring. Because of this sampling routine and because calves are being born all year round, age at sampling and sampling intervals vary for adult animals (Figure S1). Further information about the animal population can be found in Supplementary File 2.

### DNA extraction and qPCR

DNA from whole blood samples was extracted with the DNeasy Blood and Tissue spin column kit (QIAGEN) and telomere length was measured by qPCR as previously described (Seeker et al., 2016; Seeker et al., 2018; Seeker et al., 2018; Seeker, Holland, Psifidi, Banos, & Nussey, 2015). The repeatability of the assay (see Supplementary File 2 for how repeatability was calculated) was 80% and therefore delivers robust results for further interpretation (Nettle, Seeker, Nussey, Froy, & Bateson, 2019). A full description of our DNA extraction and qPCR protocols including quality control steps can also be found in Supplementary File 2.

### Statistical analysis

We have shown before that our RLTL data are significantly affected by qPCR plate and qPCR row (Seeker et al., 2016; Seeker et al., 2018). To account for those known sources of measurement error, we used the residuals of a linear model that corrected all RLTL measurements for qPCR plate and row, by fitting plate and row as fixed factors in the model. These residual RLTL measures were used in all subsequent calculations and models of telomere dynamics. RLTL change was calculated as the difference between two subsequent adjusted RLTL measurements within individuals (RLTL change = RLTLt - RLTLt-1). To investigate which factors affect the direction and amount of RLTL change, a linear mixed model was fitted with RLTL change as response variable and animal identity as random effect. The following factors were included as fixed effects in the model to test their relationship with RLTL change: genetic line, feed group and birth year of the animal, age at sampling (at time t), and the occurrence of a health event within two weeks before or after sampling (at time t). The time difference between consecutive samplings in days was fitted as a covariate. Non-significant fixed effects (p>0.05) were backwards eliminated from the model. Age at sampling was modelled as a covariate (age in years). Because we wanted to investigate if milk productivity was associated with RLTL change, the model of RLTL change was repeated for 918 change measurements of 253 animals for which milk productivity measurements were available and average lifetime milk production was fitted as additional covariate.

To investigate the association between RLTL change and productive lifespan we first focussed on RLTL change within the first year of life by only considering two RLTL measurements per animal: The first was taken shortly after birth and the second at the approximate age of one year. The change between those measurements was tested as an explanatory variable in a Cox proportional hazard model of productive lifespan.

Our data shows similar to other longitudinal studies with more than two measurements no constant attrition, maintenance or elongation, but more complex dynamics with short term changes in both directions. Example individual RLTL dynamics are shown for all cows with at least seven RLTL measurements in our dataset in figure S4. The complexity of telomere dynamics (which cannot be explained by measurement error alone) means that it is impossible to compare lifelong telomere dynamics by simply comparing slopes. To still be able to test how life-long telomere dynamics are associated with productive lifespan, we calculated three average metrics of RLTL dynamics over every individual’s lifetime: Firstly, the average of all their RLTL measures (“mean RLTL”) which investigates if animals with on average longer RLTL have a survival advantage, secondly, the average of all their RLTL change measures (“mean RLTL change”) and thirdly, the average of all their absolute RLTL change measures (“mean absolute RLTL change”). While mean RLTL change captures the direction and magnitude of RLTL changes, mean absolute RLTL change describes just the magnitude of change without considering its direction because we were interested to investigate if more change in either direction may be correlated with adverse effects. Figure S5 visualises the reasoning behind calculating these three measures of lifetime telomere length dynamics which are not surprisingly moderately correlated with one another (r ranged from −0.53 to 0.30, Figure S6). We used Cox proportional hazard models of productive lifespan that also included mean milk production as covariate and included mean RLTL, mean RLTL change and mean absolute RLTL change as explanatory variables first separately and then together to test their association with productive lifespan first separately, then while accounting for the effect of the other two measures. We have reported before that on average RLTL shortens in this population of dairy cattle within the first year of life while remaining relatively stable thereafter at the population level (Seeker et al., 2018; Seeker, et al., 2018). We were concerned that the initial RLTL attrition may contribute proportionally more to overall RLTL shortening in short-lived animals with less follow up samples which may explain a correlation of more telomere attrition with a shorter productive lifespan. To ensure the relationship was not entirely driven by this correlation, we tested all models again while excluding all measurements that were taken shortly after birth and then also while excluding all animals with less than three RLTL measurements. Lastly, we tested if the previously reported effect of RLTL at the age of one year on productive lifespan (Seeker, Ilska, Psifidi, Wilbourn, Underwood, Fairlie, Holland, Froy, Salvo-Chirnside, et al., 2018) remained statistically significant when tested in a Cox proportional hazard model together with milk productivity and mean RLTL change.

All statistical analyses were performed in R studio with R 3.1.3. (R Core Team, 2014). Mixed-effects models were implemented using the ‘lme4’ library (Bates, Mächler, Bolker, & Walker, 2015), while Cox proportional hazard models were implemented using the library survival (Therneau, 2015) and figures were generated with the library ‘ggplot2’ (Wickham, 2009).

## Results

We used blood samples collected between 2008 and 2014 to measure relative leukocyte telomere length (RLTL) by monoplex qPCR in 1,328 samples from 308 individuals. On average 4.3 (range: 2-8) telomere measurements were made of each individual, including the first measurement within 15 days of birth and a variable number of subsequent measures (Figure S5 A). RLTL measures, adjusted for qPCR plate and row (see Methods), were approximately normally distributed (Figure S7 B). From these longitudinal data we calculated 1,020 measures of change in adjusted RLTL across subsequent measurements of individual animals. RLTL change also showed approximately a normal distribution with a mean statistically significantly smaller than zero (Figure S7 C; t = −5.844, df = 1019, p-value <0.001) indicating that telomere shortening overall predominated. Animals varied in the amount and direction of RLTL change across consecutive measurements, with a fairly even proportion of individuals increasing (43.2%) and decreasing (56.8%) in RLTL over time. We next ran a series of models to test whether known individual, genetic and environmental variables could explain variation in RLTL change. Only age at sampling was significant in these models; genetic group, feed group, birth year, the time difference between samplings in days, and the occurrence of a health event within two weeks of sampling were not significant (Table S1). Also, average lifetime milk productivity was not statistically associated with RLTL change, when tested in a subset of animals (253 animals with 918 RLTL change measurements) that had milk productivity measurements available (Table S2). We have previously shown that, on average across this population, RLTL declined over the first year of life but showed no systematic change with age thereafter (Seeker, Ilska, Psifidi, Wilbourn, Underwood, Fairlie, Holland, Froy, Salvo-Chirnside, et al., 2018; Seeker, Ilska, Psifidi, Wilbourn, Underwood, Fairlie, Holland, Froy, Bagnall, et al., 2018). Consistent with this, we found that average RLTL change across consecutive measurements was only significantly negative (indicating a tendency for attrition over time) when the first measurement was made close to birth and the follow up measurement at the age of around 1 year (Figure 1A, estimate= −0.115, SE= 0.01, t= −11.512, p<0.001; Table S3). Consecutive RLTL measurements made on the same individual were overall moderately positively and significantly correlated (r=0.38, p<0.001; Figure 1B), supporting our previously reported moderate and significant individual repeatability of RLTL (Seeker, Ilska, Psifidi, Wilbourn, Underwood, Fairlie, Holland, Froy, Bagnall, et al., 2018).

**Figure 1:**
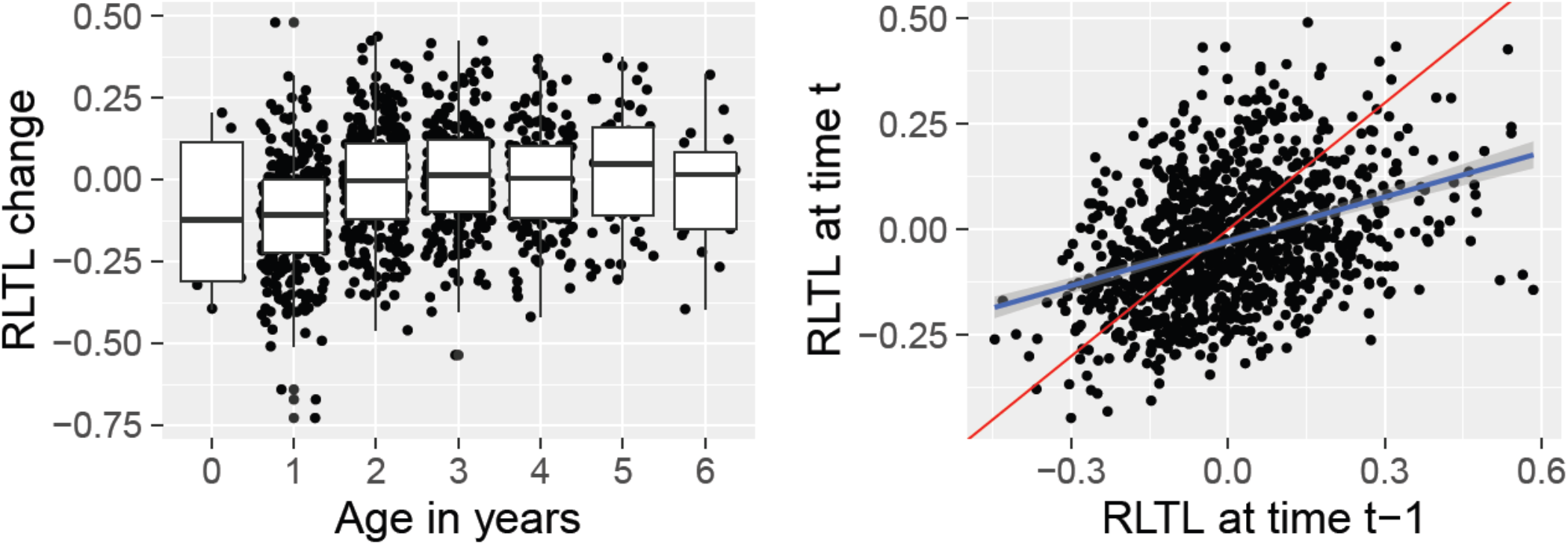
Factors that influence RLTL change. (A) Age in years at second RLTL measurement is significantly associated with average lifetime RLTL change. (B) At all measurement times the present measurement (RLTL at time t) is clearly correlated with the previous measurement (RLTL at time t-1) (estimate=0.38, p<0.001). The red line represents a perfect correlation. All RLTL measurements were pre-adjusted for qPCR plate and qPCR row to account for some of the known measurement error.

We next tested whether measured RLTL change predicted the productive lifespan of cattle. Of all dead cows (N=244) the vast majority (N=241) had survived to their first lactation, but there was considerable variation in productive lifespan beyond this point (Figure S7 D). We used a Cox proportional hazard model to test whether early life RLTL change between two samples, one taken shortly after birth and the next at an approximate age of one year, predicted productive lifespan and found that greater early life RLTL attrition was associated with a shorter productive lifespan (N= 291, coefficient= −1.141, SE= 0.391, p= 0.004, Figure 2, Table S4).

**Figure 2:**
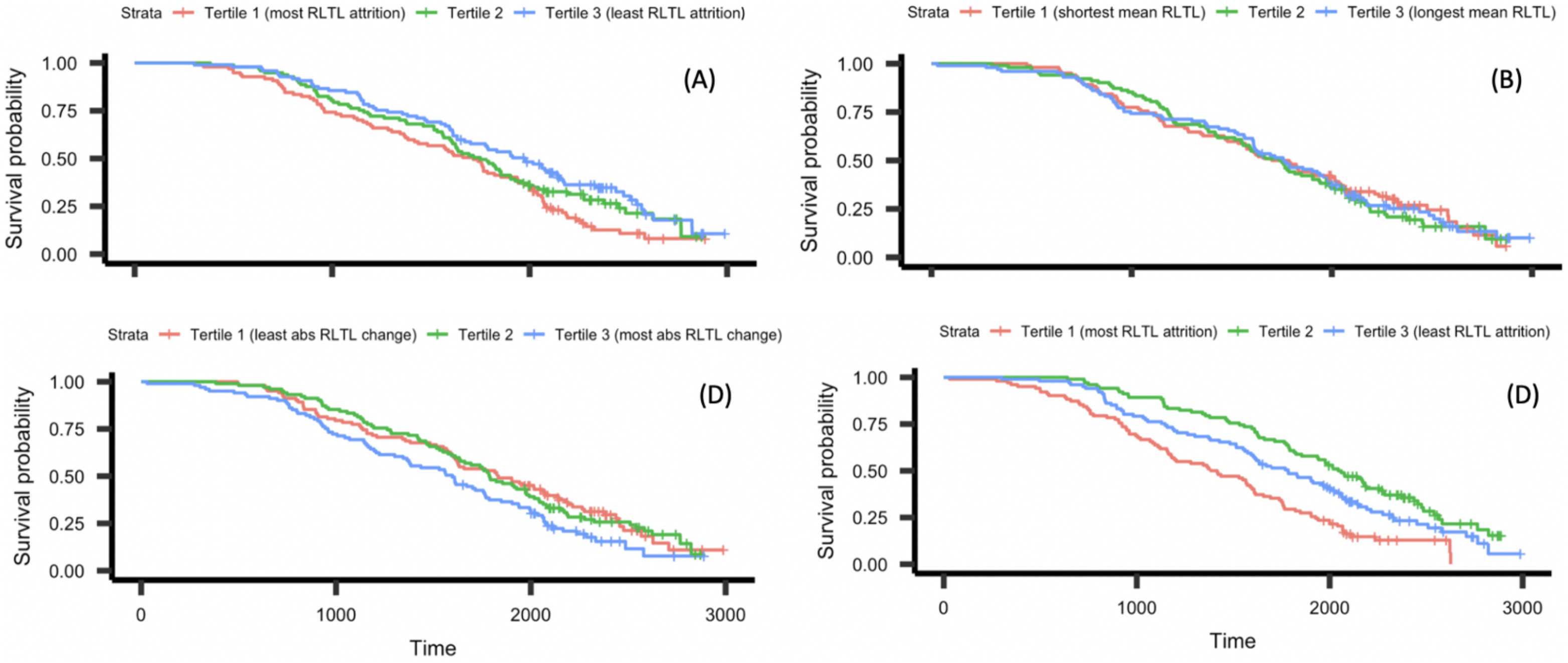
Kaplan-Meier curves for cattle divided into tertiles based on different telomere length measures over their lifetimes, illustrating relationships between productive lifespan and lifelong telomere dynamics. (A) More RLTL attrition within the first year of life was associated with a shorter productive lifespan (N= 291, coefficient = −0.23, SE= 0.08, Wald test= 7.59 on 1df, p= 0.006). (B) Lifetime mean RLTL was not significantly associated with productive lifespan (N=305, coefficient= 0.01, SE=0.08, Wald test= 0.03 on 1 df, p= 0.9). (C) Greater mean absolute RLTL change measured over the lifetime was associated with shorter productive lifespan (N=305, coefficient=0.18, SE=0.08, Wald test= 4.1 on 1 df, p=0.03). (D) Greater mean lifetime RLTL attrition was associated with shorter productive lifespan (N= 305, estimate=-0.25, SE=0.09, Wald test= 8.85 on 1 df, p=0.003).

When each measure of lifetime RLTL dynamics was tested in a separate cox proportional hazard model of productive lifespan, mean RLTL did not significantly predict productive lifespan (coefficient= 0.341, SE= 0.591, p=0.564, Figure 2B, Table S4), but both mean RLTL change (coefficient= −5.209, SE= 0.845, p<0.001; Figure 2D, Table S4) and mean absolute RLTL change (coefficient= 2.939, SE= 0.970, p=0.002; Figure 2C, Table S4) were significantly associated with productive lifespan. When all three measures of lifetime RLTL dynamics were included in the same model only mean RLTL change remained significant (coefficient = −4.758, SE= 1.018, p<0.001; Table S4). This implies that the relationship between productive lifespan and mean absolute RLTL change was largely due to covariance with mean RLTL change. Thus, individuals that experienced greater telomere attrition over their lifetimes had a shorter productive lifespan and direction of RLTL change (rather than simply absolute magnitude) was an important aspect of this relationship. To visualise the relationship between RLTL change measurements and productive lifespan using Kaplan-Meier plots (Figure 2), continuous RLTL measures were transformed to a discrete scale by grouping them into tertiles (Figure S8). Cox proportional hazard models based on these tertile groupings of RLTL measures showed similar results to those reported above (Figure 2).

The relationship between mean RLTL change and productive lifespan was robust to the inclusion of milk production (a physiological stressor, which is positively associated with productive lifespan as cows with a low milk yield are more likely to be culled at a younger age) in the model (Table S5). If RLTL declines most within the first year of life (as Figure 1A indicates), the association between change in RLTL and productive lifespan may be driven by the fact that this initial decline contributes relatively more to estimates of mean RLTL change in shorter-lived individuals than in animals that have more follow up samples with more moderate change measures available. We therefore repeated the analysis excluding the early life RLTL measurements and found that mean RLTL change still predicted productive lifespan (N=253, coefficient= −5.056, SE= 1.315, p<0.001, Table S6). The relationship between mean RLTL change and productive lifespan also remained statistically significant, when animals with less than 3 samples were excluded from the analysis (N= 213, coefficient= −3.47, SE= 1.34, Wald test = 6.7 on 1 df, p= 0.01) and when RLTL at 1 year of age (which we previously found to predict productive lifespan (Seeker, Ilska, Psifidi, Wilbourn, Underwood, Fairlie, Holland, Froy, Salvo-Chirnside, et al., 2018)) was included in the models (Table S7).

## Discussion

We have presented, to our knowledge, the first demonstration in any vertebrate that lifelong variation in telomere attrition rather than variation in constitutive individual differences in average telomere length predict health outcomes and lifespan. While there is mounting evidence that telomere length predicts mortality, health and life history in humans as well as birds and non-human mammals (Bakaysa et al., 2007; Bichet et al., 2019; Boonekamp et al., 2013; Cawthon, Smith, O’Brien, Sivatchenko, & Kerber, 2003; Fairlie et al., 2015; Mark Haussmann, Winkler, & Vleck, 2005; Heidinger et al., 2012; Nettle, Monaghan, Boner, Gillespie, & Bateson, 2013; Voillemot et al., 2012; Wilbourn et al., 2018), very few studies have been able to accumulate long-term longitudinal data capable of differentiating the role of among- and within-individual variation in TL to such relationships. Our data support the contention that within-individual directional change over time in TL is more important than stable among-individual differences in predicting overall health. Individuals in our study which showed a greater propensity to shorten RLTL across sampling points were culled earlier, mainly due to health issues described above. There was no relationship between productive lifespan and an individual’s average RLTL. Future studies will show how well our results generalise to other systems as telomere biology is variable amongst species. Cattle telomere biology seems to be similar to other ruminants, horses, zebras, tapirs some whales and primates including humans in that they have relatively short telomeres and a tight regulation of telomerase expression (Gomes et al., 2011). If our results generalise to some of those other systems and contexts, they have important implications for the utility of TL as a biomarker of health and fitness, lending support to the idea that TL change is an indirect marker reflecting past physiological insults and stress rather than an indicator of constitutive or genetically-based robustness to life’s challenges. Our data also highlight the importance of collecting longitudinal telomere measurements, by showing that in some species it is within-individual change over time in TL that carries the important biological signal.

Our study animals varied considerably in the magnitude and direction of RLTL change across consecutive sampling points. This is in accordance with other longitudinal studies that have reported a wide variation in TL change, alongside evidence that a large proportion of individuals show telomere lengthening over time (Bize et al., 2009; Chen et al., 2011; Farzaneh-Far et al., 2010; Gardner et al., 2005; Huzen et al., 2014; Kark, Goldberger, Kimura, Sinnreich, & Aviv, 2012; Nordfjäll et al., 2009; Shalev et al., 2012; Steenstrup et al., 2013; Svenson et al., 2011). Whilst the vast majority of studies linking telomere length with health and fitness remain cross-sectional, a growing number of longitudinal studies have found that individuals that lose more TL across samples have reduced survival prospects (Barrett et al., 2013; Bize et al., 2009; Boonekamp et al., 2014; Goglin et al., 2016). However, these studies tend to involve two samples taken per individual or a follow-up period that represents only a small fraction of the total lifespan of the study species. A lifelong study of TL and lifespan in zebra finches found that TL measured in early life was a better predictor of subsequent survival than measures from later life, but the role of change in telomere length across sampling points was not investigated (Heidinger et al., 2012). Here we show that, despite TL being moderately consistent across the lifetimes of individuals, there remains considerable within-individual variation and the pattern of change in TL over an individual’s life is highly dynamic. Short-term environmental fluctuations impacting TL dynamics could be responsible and may impact individuals in different ways. In humans many lifestyle factors such as socio-economic status, psychological stress, smoking and body mass index are known to influence telomere length (Cherkas et al., 2008; Epel, Daubenmier, Moskowitz, Folkman, & Blackburn, 2009; Huzen et al., 2014; Kim et al., 2009; Shalev et al., 2012). In animal studies chronic infection and exposure to stressful situations have been linked to telomere shortening (Asghar, Hasselquist, Zehtindjiev, Westerdahl, & Bensch, 2015; Nettle et al., 2013; Reichert et al., 2014). Whilst we were unable to identify variables other than age which predicted within-individual TL change in our study system, individuals with a greater propensity to lose TL over time had shorter productive lives, implying changes in TL reflect important environmental or physiological variation linked to health. From a telomere biology perspective, it would be interesting to investigate changes not only in mean TL but also in its variance over time, preferentially using a technique like STELA or TESLA that allows the detection of critically short telomeres as well (Baird, Rowson, Wynford-Thomas, & Kipling, 2003; Lai et al., 2017). However, if the focus is to find good markers for animal breeding and health those techniques are not economically practical.

Our results have important implications for the application of TL as a biomarker within animal breeding and health. Productive lifespan describes the time from birth to culling and differs from measures of “longevity” or “lifespan” in human or wild animal studies because the animals are culled for reasons relating mainly to health and productivity. From an animal production perspective, a biomarker which could predict the productive lifespan of livestock from a relatively early age would represent an exciting target phenotype for use in selective breeding if heritable or for use in animal welfare monitoring if influenced by the environment. In the present study we showed that RLTL attrition predicted productive lifespan. The fact that both early life and lifelong RLTL shortening predicts earlier culling does support recent suggestions that TL may be useful as a biomarker of stress and welfare in livestock (Melissa Bateson, 2016; Brown, Dechow, Liu, Harvatine, & Ott, 2012). Many previous studies in birds and humans have found that experience of past stress is associated with shorter telomeres (Angelier, Vleck, Holberton, & Marra, 2013; Asghar et al., 2015; Cherkas et al., 2006; Entringer et al., 2011; Entringer et al., 2013; Epel et al., 2004; Haussmann, Longenecker, Marchetto, Juliano, & Bowden, 2012; Nettle et al., 2017). In our study, cattle showing RLTL attrition may be those that have experienced some form of recent immediate stress or minor infection. Repeated bouts of such stress, reflected by higher average RLTL attrition through life, may result in the development of health issues that lead to culling. Whilst the results of the present study suggest that longitudinal RLTL change itself may be a useful biomarker of health and welfare in livestock, a key area for further study is the causes of variation in RLTL change, as these may help us to understand sources of physiological stress for domestic animals and help improve animal welfare and productivity.

## Acknowledgements

We thank everyone involved in collecting blood samples on the Crichton farm, particularly Hannah Dykes and Isla McCubbin. We also thank the SynthSys facility at the University of Edinburgh, particularly Eliane Salvo-Chirnside, for their support with the liquid handling robot. Our work was funded by the Biotechnology and Biological Sciences Research Council (BB/L007312/1; http://www.bbsrc.ac.uk/) awarded to GB. LS received a studentship from Scotland’s Rural College (http://www.sruc.ac.uk/).

## Data Accessibility

The complete dataset will be accessible after acceptance on GitHub.

## Author contributions

All authors designed the experiment together. LS, SU, RW, JF, RH and AB collected the blood samples and processed them in the lab. LS analysed the data with help from HF, JI, AP, GB and DN. LS wrote the initial draft of the manuscript, DN and GB edited. All authors were involved in revising the manuscript. BW, MC, GB and DN obtained funding for the project.

## Supplementary File 1

### Supplementary Figures and Tables

#### Supplementary Figures

**Figure S1:**
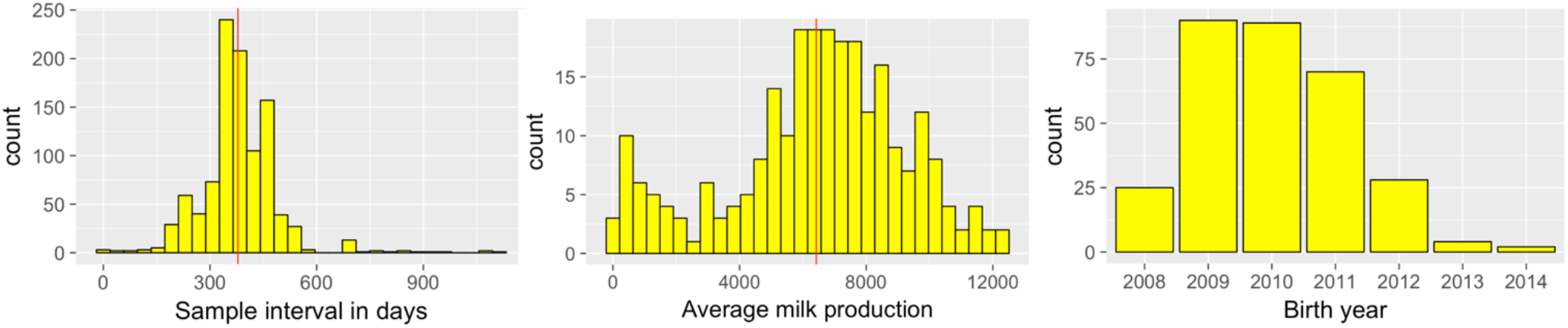
Left: Sample interval between two consecutive samples varied considerably with a mean of approximately one year. Middle: Average lifetime milk production for all animals that started at least the first lactation. Right: Number of animals born in specific years between 2008 and 2014.

**Figure S2:**
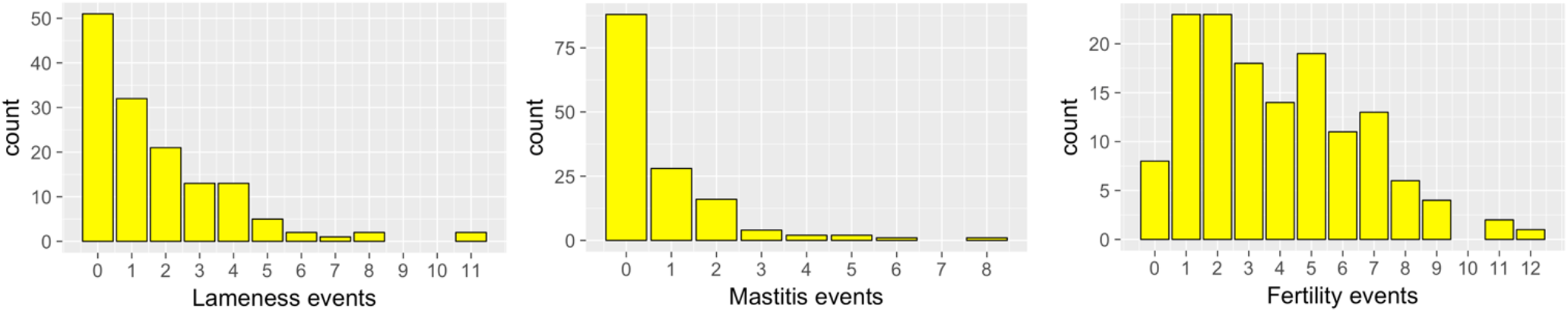
Number of specific health events per animal.

**Figure S3:**
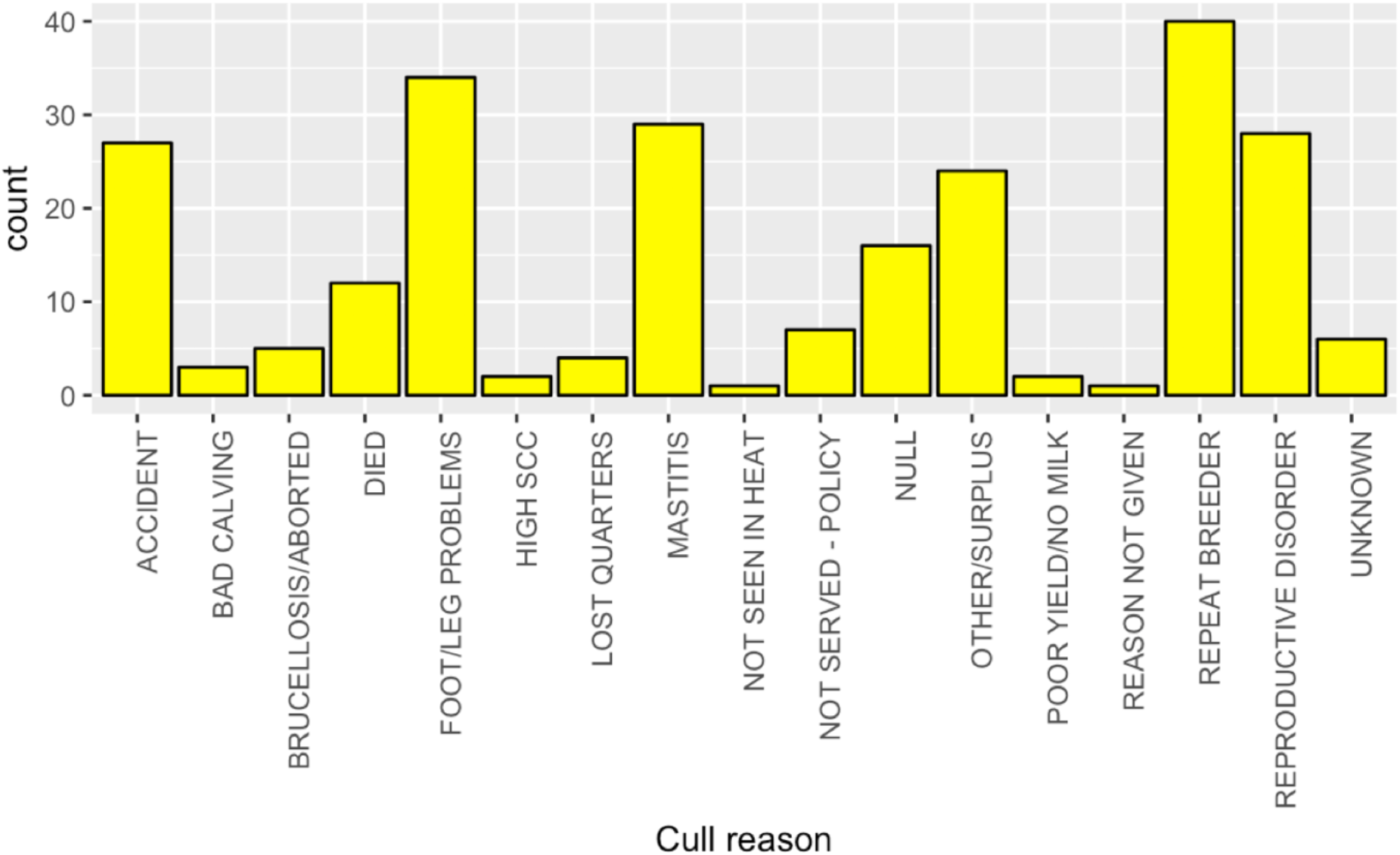
Reasons for culling. High SCC: High somatic cell count as a marker for (subclinical) mastitis. Repeat breeder is a cow that fails to conceive after multiple inseminations.

**Figure S4:**
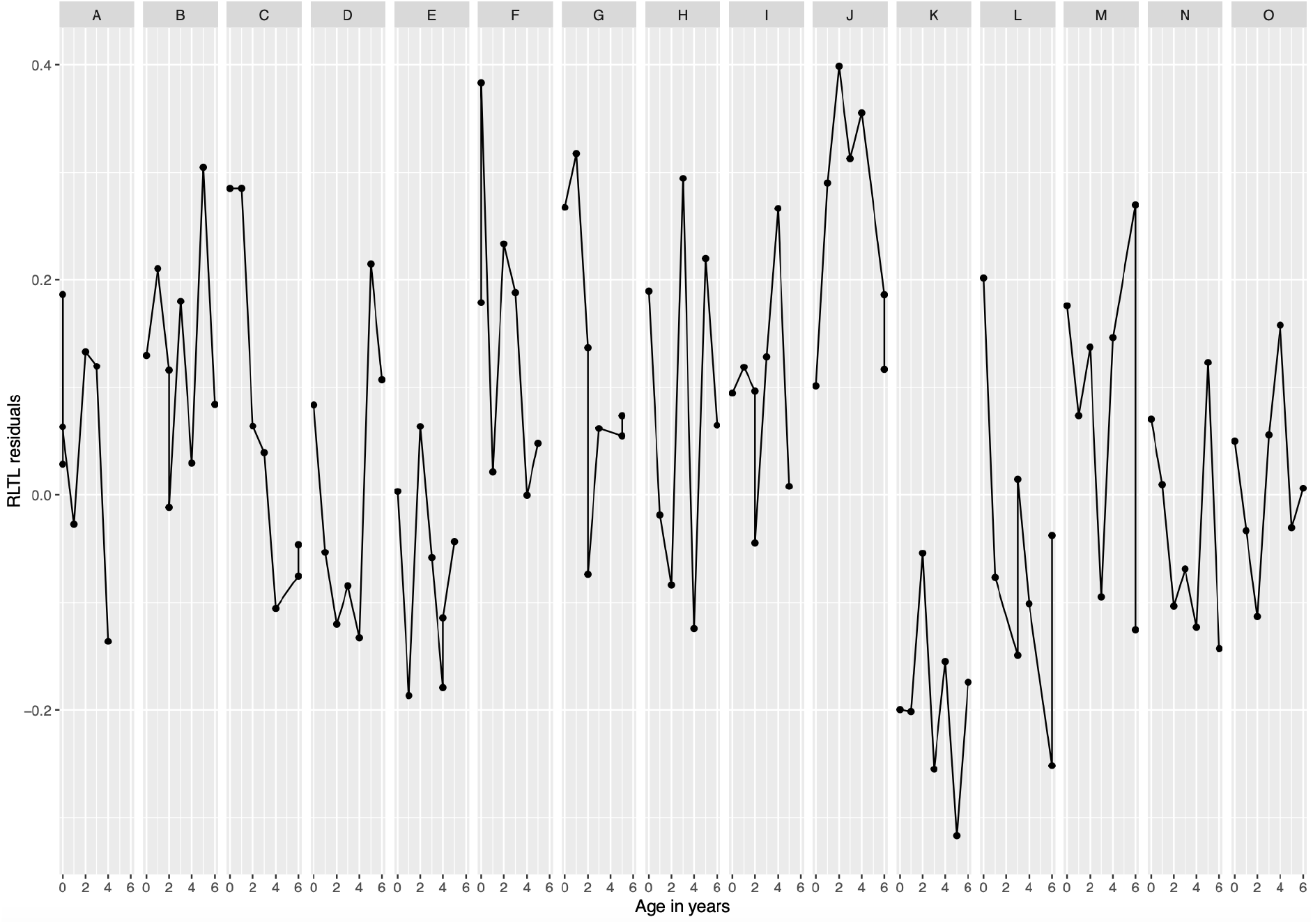
Relative leukocyte telomere length (RLTL) dynamics is variable amongst individuals. RLTL corrected for qPCR plate and qPCR row (see methods) over age in years for individual animals (A-O) that had at least seven telomere length measurements present.

**Figure S5:**
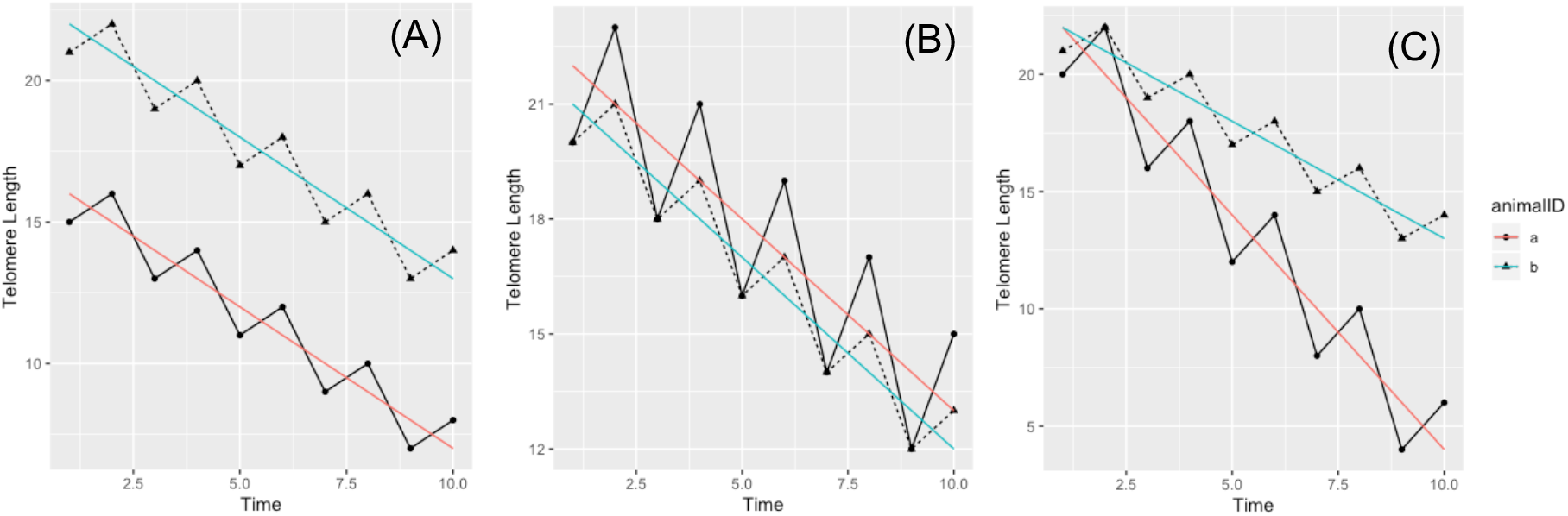
Visualisation of reasoning behind the calculation of different life-long telomere change measures. Hypothetical scenario (A): differences in mean relative leukocyte telomere length (RLTL). Animals differ in their mean telomere length while telomere attrition rate and variance in telomere length are similar. It is suspected that animals with longer mean RLTLs over life (like animal b) have a survival advantage over those with on average shorter RLTLs (like animal a). Hypothetical scenario (B): Differences in mean absolute RLTL change. Animals do not differ significantly in their telomere length or their overall attrition rate. However, they show a difference in their magnitude of short-term telomere change. It is hypothesised that animals that are able to better maintain their telomere length and show lees absolute telomere change (like animal b) have a survival advantage over those that show extreme telomere change in both directions (like animal a). Hypothetical scenario (C): Differences in mean RLTL change. Some animals show a faster overall telomere attrition (like animal a) compared to others (like animal b) and it is hypothesised that fast telomere attrition is associated with a shorter lifespan.

**Figure S6:**
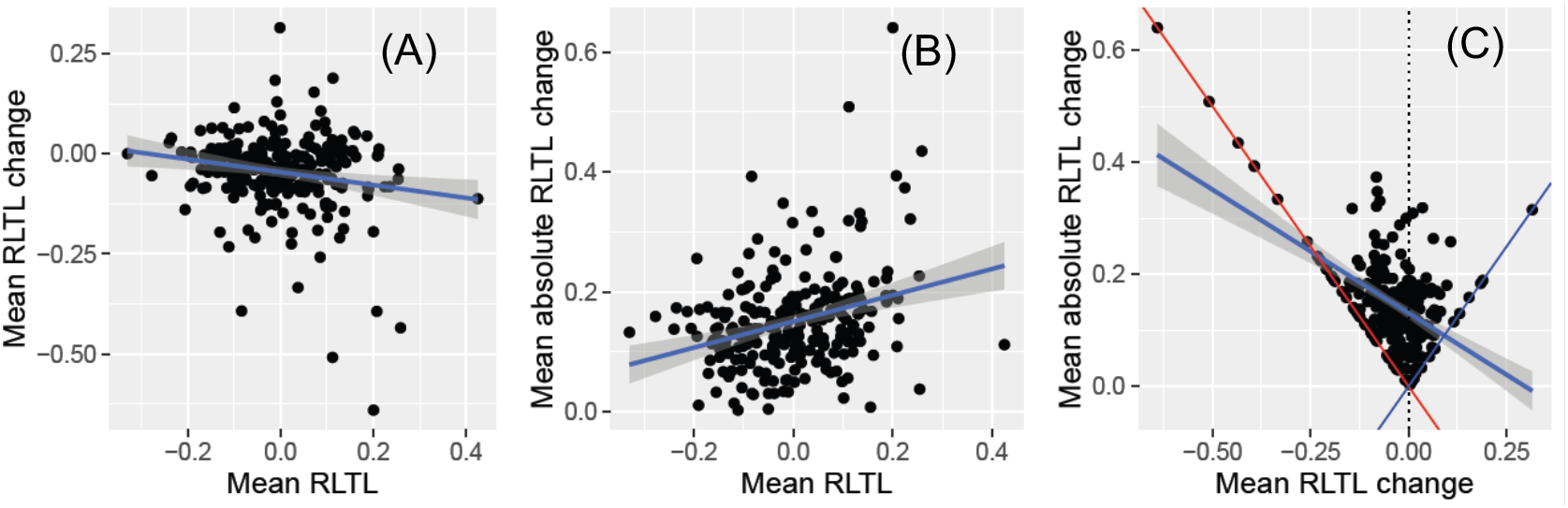
Correlations between three different measures of lifetime relative leukocyte telomere length (RLTL) dynamics calculated for 244 dead animals. (A) Mean RLTL and mean RLTL change: r=-0.18, p=0.004; (B) Mean RLTL and mean absolute RLTL change: r=0.30, p<0.001. (C) Mean RLTL change and mean absolute RLTL change: r=-0.53, p<0.001. The red line represents perfect negative correlation, blue line represents perfect positive correlation. Dotted line represents data that shows a variable amount of absolute change that cancels each other out to result in no overall RLTL change.

**Figure S7:**
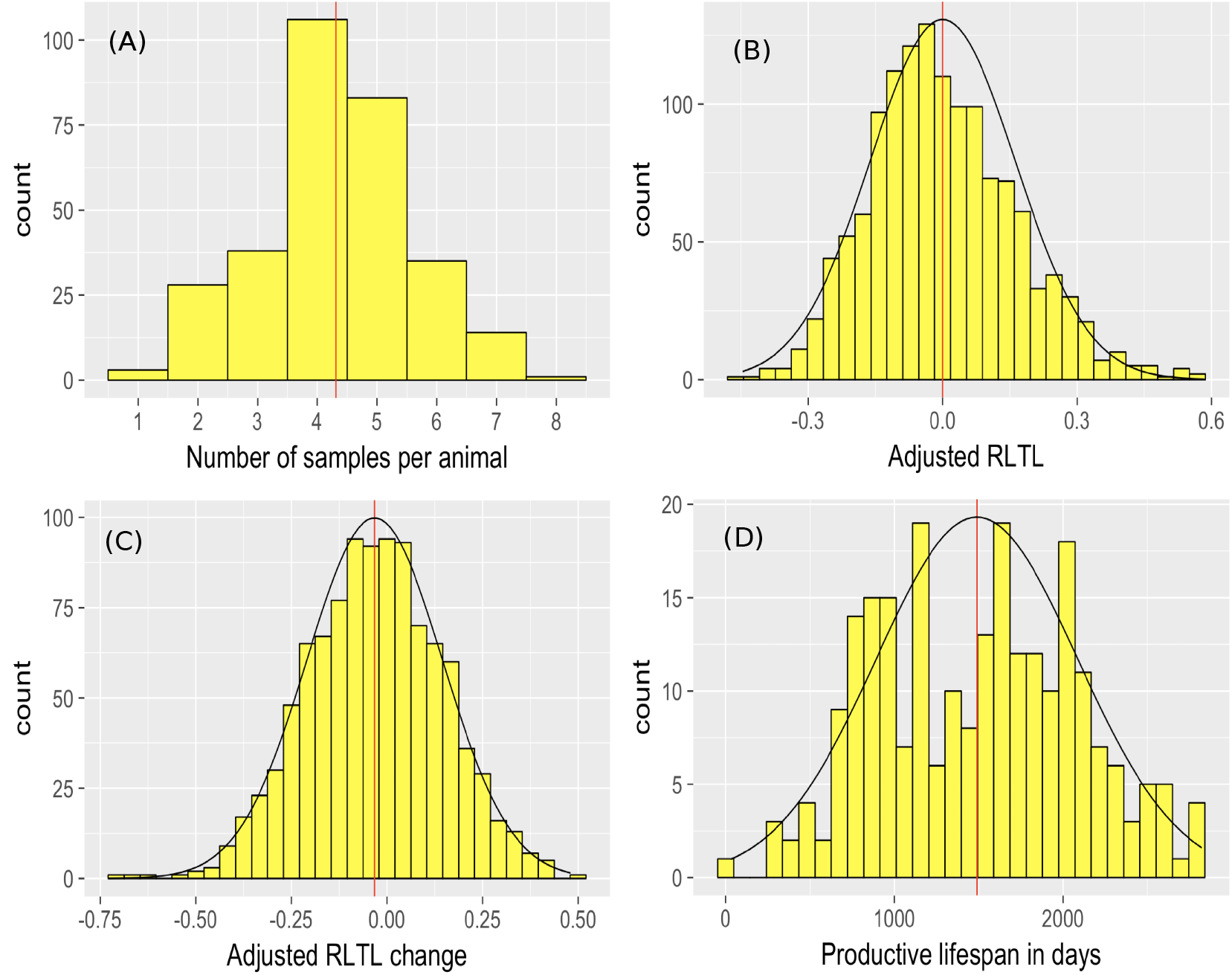
Histograms of (A) Number of RLTL measurements per animal, (B) RLTL residuals after correcting for qPCR plate and row, (C) measurement to measurement change in RLTL residuals (after correcting for qPCR plate and row) and (D) productive lifespan in days; Red lines indicate mean.

**Figure S8:**
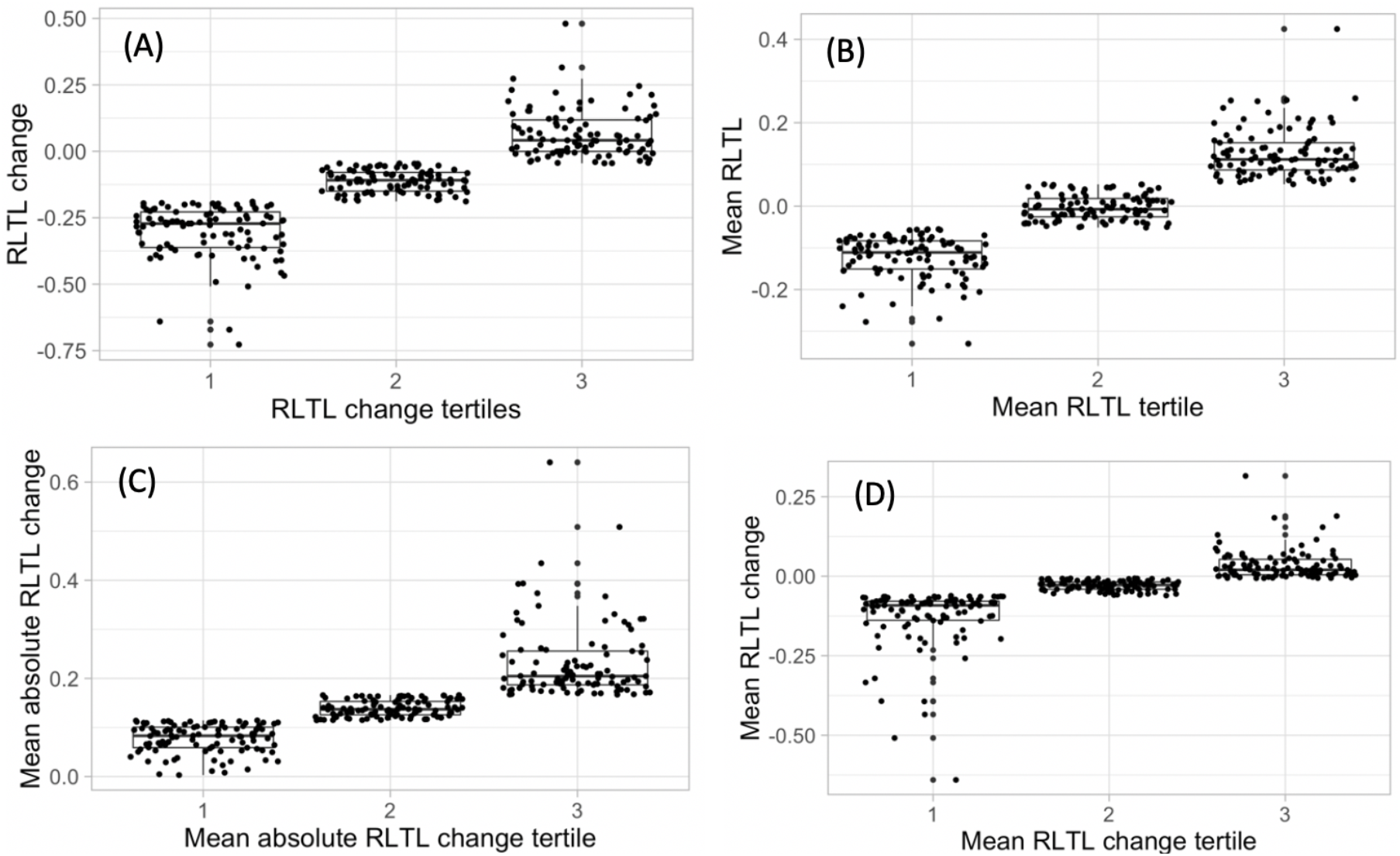
Grouping of RLTL measurements into tertiles. (A) RLTL change within the first year of life, (B) mean RLTL, (C) mean absolute RLTL change, (D) mean RLTL change.

#### Supplementary Tables

**Table S1:**
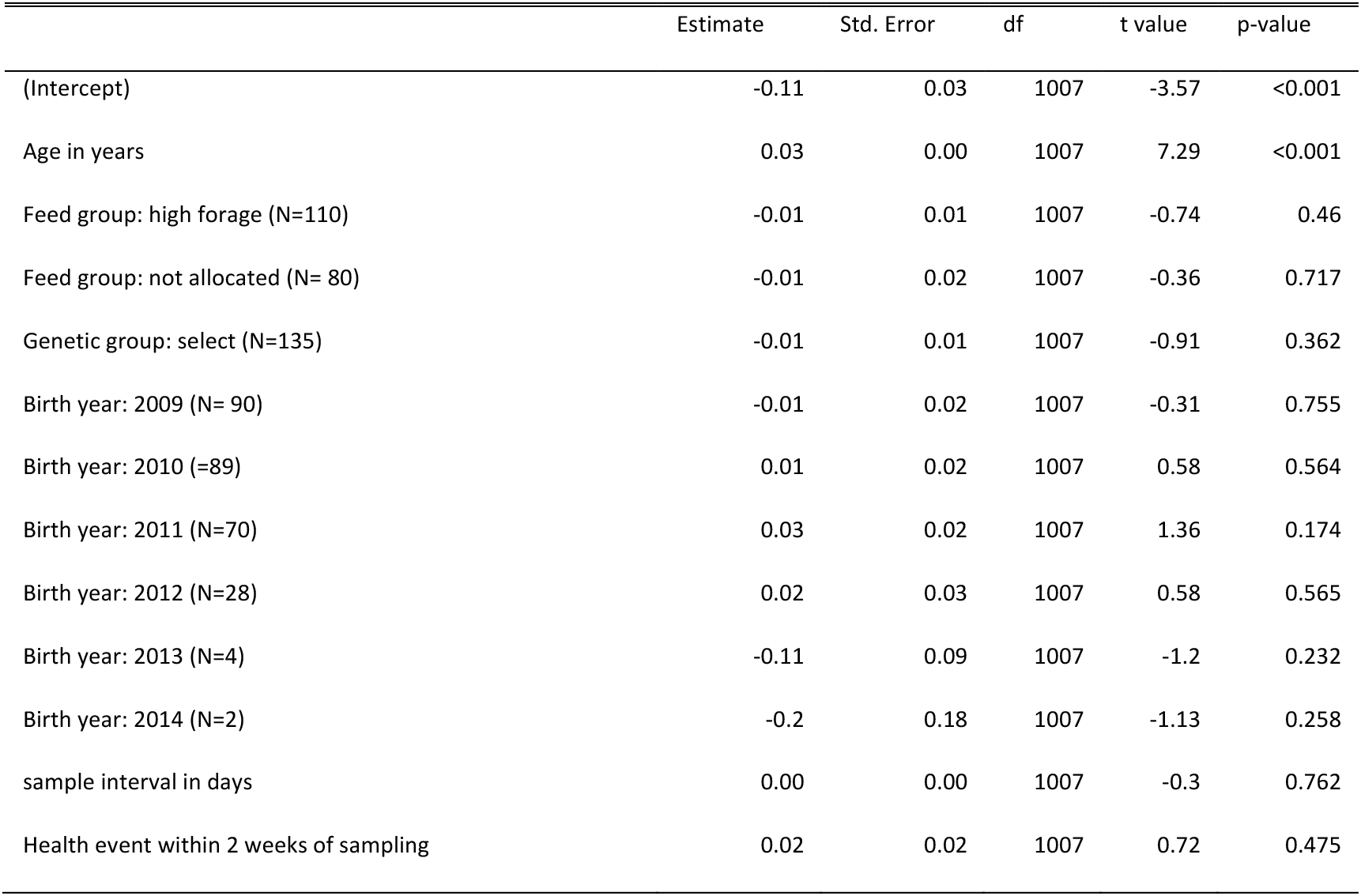
Effect sizes and significance of fixed effects from linear mixed-effects model of RLTL change, adjusted for qPCR plate and row with animal identity fitted as a random effect.

|  | Estimate | Std. Error | df | t value | p-value |
| --- | --- | --- | --- | --- | --- |
| (Intercept) | -0.11 | 0.03 | 1007 | -3.57 | <0.001 |
| Age in years | 0.03 | 0.00 | 1007 | 7.29 | <0.001 |
| Feed group: high forage (N=110) | -0.01 | 0.01 | 1007 | -0.74 | 0.46 |
| Feed group: not allocated (N= 80) | -0.01 | 0.02 | 1007 | -0.36 | 0.717 |
| Genetic group: select (N=135) | -0.01 | 0.01 | 1007 | -0.91 | 0.362 |
| Birth year: 2009 (N= 90) | -0.01 | 0.02 | 1007 | -0.31 | 0.755 |
| Birth year: 2010 (=89) | 0.01 | 0.02 | 1007 | 0.58 | 0.564 |
| Birth year: 2011 (N=70) | 0.03 | 0.02 | 1007 | 1.36 | 0.174 |
| Birth year: 2012 (N=28) | 0.02 | 0.03 | 1007 | 0.58 | 0.565 |
| Birth year: 2013 (N=4) | -0.11 | 0.09 | 1007 | -1.2 | 0.232 |
| Birth year: 2014 (N=2) | -0.2 | 0.18 | 1007 | -1.13 | 0.258 |
| sample interval in days | 0.00 | 0.00 | 1007 | -0.3 | 0.762 |
| Health event within 2 weeks of sampling | 0.02 | 0.02 | 1007 | 0.72 | 0.475 |

**Table S2:** Effect sizes and significance of fixed effects from linear mixed-effects model of RLTL change, adjusted for qPCR plate and row with animal identity fitted as a random effect, including average lifetime milk production. Model identical to that presented in Table S1, except for inclusion of this additional fixed effect. Number of animals:253, number of samples: 918)

|  | Estimate | Std. Error | df | t value | p-value |
| --- | --- | --- | --- | --- | --- |
| (Intercept) | -0.12 | 0.04 | 906 | -3.28 | 0.001 |
| Age in years | 0.03 | 0.01 | 906 | 6.76 | <0.001 |
| Feed group: High fibre (N=108) | 0.00 | 0.01 | 906 | -0.25 | 0.802 |
| Feed group: Not allocated (N= 29) | 0.01 | 0.03 | 906 | 0.54 | 0.59 |
| Genetic group: select (N=117) | -0.01 | 0.01 | 906 | -0.94 | 0.35 |
| Birth year: 2009 (N=77) | -0.01 | 0.02 | 906 | -0.47 | 0.639 |
| Birth year: 2010 (N=79) | 0.01 | 0.02 | 906 | 0.43 | 0.666 |
| Birth year: 2011 (N=62) | 0.03 | 0.02 | 906 | 1.21 | 0.226 |
| Birth year: 2012 (N=10) | 0.03 | 0.04 | 906 | 0.72 | 0.472 |
| sample interval in days | 0.00 | 0.00 | 906 | -0.63 | 0.529 |
| Health event within 2 weeks of sampling | 0.02 | 0.02 | 906 | 0.75 | 0.452 |
| Average milk productivity | 0.00 | 0.00 | 906 | 1.12 | 0.261 |

**Table S3:**
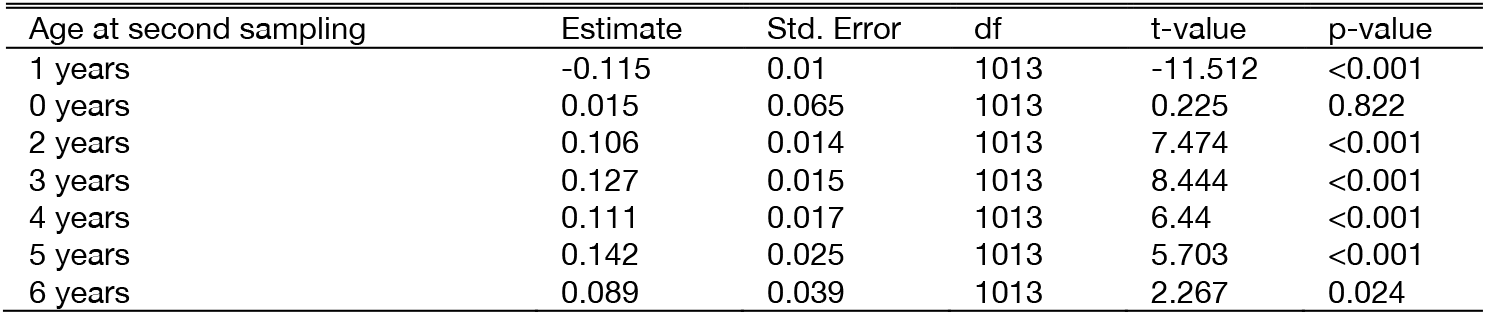
Effect sizes and significance of fixed effects of age at second sampling from linear mixed-effects model of RLTL change, adjusted for qPCR plate and row with animal identity fitted as a random effect, Age at second sampling in years was tested as a factor to illustrate that average RLTL change across consecutive measurements was only significantly negative (indicating a tendency for attrition over time) when the first measurement was made close to birth and the follow up measurement at the age of around 1 year.

| Age at second sampling | Estimate | Std. Error | df | t-value | p-value |
| --- | --- | --- | --- | --- | --- |
| 1 years | -0.115 | 0.01 | 1013 | -11.512 | <0.001 |
| 0 years | 0.015 | 0.065 | 1013 | 0.225 | 0.822 |
| 2 years | 0.106 | 0.014 | 1013 | 7.474 | <0.001 |
| 3 years | 0.127 | 0.015 | 1013 | 8.444 | <0.001 |
| 4 years | 0.111 | 0.017 | 1013 | 6.44 | <0.001 |
| 5 years | 0.142 | 0.025 | 1013 | 5.703 | <0.001 |
| 6 years | 0.089 | 0.039 | 1013 | 2.267 | 0.024 |

**Table S4:** Results of cox proportional hazards models testing association of relative leukocyte telomere length (RLTL) change measurements (on continuous scales) with productive lifespan. SE= standard error, CI = confidence interval

| RLTL measure | Coefficient (SE) | Hazard ratio | 95% CI (hazard ratio) | exp(-Coefficient) | z | p-value |
| --- | --- | --- | --- | --- | --- | --- |
| **1) RLTL change within the first year of life (N=291)** |  |  |  |  |  |  |
| RLTL change | -1.141 (0.391) | 0.320 | 0.148 -0.688 | 3.129 | -2.914 | 0.004 |
| **1) Each RLTL change measurements is tested in separate models (N=305)** |  |  |  |  |  |  |
| Mean RLTL | 0.341 (0.591) | 1.406 | 0.442-4.473 | 0.711 | 0.577 | 0.564 |
| Mean RLTL change | -5.209 (0.845) | 0.005 | 0.001-0.029 | 183 | -6.166 | <0.001 |
| Mean absolute RLTL change | 2.939 (0.970) | 18.8982 | 2.824-126.5 | 0.053 | 3.031 | 0.002 |
| **2) All RLTL change measurements are tested in the same model (N=305)** |  |  |  |  |  |  |
| Mean RLTL | 0.053 (0.606) | 1.054 | 0.321-3.456 | 0.9488 | 0.087 | 0.931 |
| Mean RLTL change | -4.758 (1.018) | 0.009 | 0.001-0.063 | 116.722 | -4.676 | <0.001 |
| Mean absolute RLTL change | 0.776 (1.076) | 2.173 | 0.264-17.910 | 0.460 | 0.721 | 0.471 |

**Table S5:** Results of cox proportional hazards models testing association of relative leukocyte telomere length (RLTL) change measurements (on continuous scales) with productive lifespan, including measures of average lifetime milk production. Average lifetime milk production was tested in the same models to account for the fact that cows may be culled because of poor productivity. SE= standard error, CI = confidence interval

| Factor | Coefficient (SE) | Hazard ratio | 95% CI (hazard ratio) | exp(-Coefficient) | z | p-value |
| --- | --- | --- | --- | --- | --- | --- |
| **1) Mean RLTL, N=253** |  |  |  |  |  |  |
| Mean RLTL | 0.232 (0.671) | 1.260 | 0.339-4.693 | 0.793 | 0.345 | 0.730 |
| Average lifetime milk production | -9.933 *10<sup>-5</sup> (2.822 *10<sup>-5</sup>) | 1.000 | 0.9998-1.000 | 1.0001 | -3.520 | <0.001 |
| **2) Mean RLTL change, N=253** |  |  |  |  |  |  |
| Mean RLTL change | -5.035 (1.315) | 0.007 | 0.001-0.086 | 153.7 | -3.829 | <0.001 |
| Average lifetime milk production | -9.057 *10<sup>-5</sup> (2.819 *10<sup>-5</sup>) | 1.000 | 0.0005-0.086 | 1.000 | -3.212 | 0.001 |
| **3) Mean absolute RLTL change, N=253** |  |  |  |  |  |  |
| Mean absolute RLTL change | 2.411 (1.146) | 11.15 | 1.1793-105.4 | 0.090 | 2.104 | 0.035 |
| Average lifetime milk production | -1.018 *10<sup>-4</sup> (2.818 *10<sup>-5</sup>) | 1.000 | 1.000-1.000 | 1.000 | -3.611 | <0.001 |
| **4) All telomere change measures tested in the same model, N=253** |  |  |  |  |  |  |
| Mean RLTL | -0.070 | 0.933 | 0.244-3.564 | 1.072 | -0.102 | 0.919 |
| Mean RLTL change | -4.525 (1.387) | 0.011 | 0.001-0.164 | 92.303 | -3.262 | 0.001 |
| Mean absolute RLTL change | 1.237 (1.224) | 3.447 | 0.313-37.945 | 0.290 | 1.011 | 0.312 |
| Average lifetime milk production | -9.162 *10<sup>-5</sup> (2.824 *10<sup>-5</sup>) | 1.000 | 1.000-1.000 | 1.000 | -3.244 | 0.001 |

**Table S6:** Results of cox proportional hazards models testing association of relative leukocyte telomere length (RLTL) change measurements (on continuous scales) with productive lifespan, with first RLTL measurements (obtained close to birth) removed from analyses. N=253; SE= standard error, CI = confidence interval

| Factor | Coefficient (SE) | Hazard ratio | 95% CI (hazard ratio) | exp(-Coefficient) | z | p-value |
| --- | --- | --- | --- | --- | --- | --- |
| **1) Mean RLTL** |  |  |  |  |  |  |
| Mean RLTL | -0.485 (0.649) | 0.616 | 0.173-2.199 | 1.624 | -0.746 | 0.455 |
| Average lifetime milk production | -9.853 * 10<sup>-5</sup><br>(2.812 * 10<sup>-5</sup>) | 1.000 | 1.000-1.000 | 1.000 | -3.504 | <0.001 |
| **2) Mean RLTL change** |  |  |  |  |  |  |
| Mean RLTL change | -5.056 (1.315) | 0.006 | 0.0005-0.084 | 153.7 | -3.829 | <0.001 |
| Average lifetime milk production | -9.057* 10<sup>-5</sup><br>(2.818* 10<sup>-5</sup>) | 1.000 | 1.000-1.000 | 1.000 | -3.228 | 0.001 |
| **3) Mean absolute RLTL change** |  |  |  |  |  |  |
| Mean absolute RLTL change | 2.403 (1.147) | 11.05 | 1.168-104.6 | 0.090 | 2.095 | 0.036 |
| Average lifetime milk production | -1.018* 10<sup>-4</sup><br>(2.818* 10<sup>-5</sup>) | 1.000 | 1.000-1.000 | 1.000 | -3.610 | <0.001 |
| **4) Mean RLTL change measures tested in the same model** |  |  |  |  |  |  |
| Mean RLTL | -0.209 (0.670) | 0.812 | 0.218-3.017 | 1.232 | -0.311 | 0.756 |
| Mean RLTL change | -4.444 (1.429) | 0.012 | 0.001-0.194 | 85.107 | -3.109 | 0.002 |
| Mean absolute RLTL change | 1.281 (1.217) | 3.600 | 0.331-39.130 | 0.278 | 1.052 | 0.293 |
| Average lifetime milk production | -9.208* 10<sup>-5</sup><br>(2.820* 10<sup>-5</sup>) | 1.000 | 1.000-1.000 | 1.000 | -3.265 | 0.001 |

**Table S7:** Results of cox proportional hazard models of productive lifespan that correspond to the data shown in Figure 2. RLTL change measures were transformed to a discrete scale with 3 groups as shown in Figure S3 to allow visualisation. “Initial RLTL change” shows results for RLTL change between two measurements one shortly after birth and one at the age of 1 year and subsequent survival, whereas the other three rows show results for mean lifetime RLTL dynamics explained in the paper.

| Model | Age interval (days) | Number of animals | Number of observations (dead animals) | Number sensor ed | coef | Exp(coef) | SE(coef) | 95% interval | Exp(-coef) | z | p-value |
| --- | --- | --- | --- | --- | --- | --- | --- | --- | --- | --- | --- |
| (A) initial RLTL change | 0-3000 | 291 | 227 | 64 | -0.226 | 0.798 | 0.082 | 0.6793 - 0.9369 | 1.253 | -2.755 | 0.006 |
| (B) Mean RLTL | 0-3000 | 305 | 241 | 64 | 0.014 | 1.014 | 0.079 | 0.869 - 1.184 | 0.986 | 0.179 | 0.858 |
| (C) Mean absolute RLTL change | 0-3000 | 305 | 241 | 64 | 0.179 | 1.196 | 0.082 | 1.019 - 1.405 | 0.836 | 2.19 | 0.029 |
| (D) Mean RLTL change | 0-3000 | 305 | 241 | 64 | -0.257 | 0.773 | 0.087 | 0.652 - 0.916 | 1.294 | -2.974 | 0.003 |

**Table S8:** Results of cox proportional hazards models testing for independent associations of mean relative leukocyte telomere length (RLTL) change measurements and RLTL measured at 1 year of age on productive lifespan. Mean RLTL change remains highly statistically significant whereas RLTL at the age of 1 year becomes marginally non-significant; SE= standard error, CI = confidence interval

| Factor | Coefficient (SE) | Hazard ratio | 95% CI (hazard ratio) | exp(-Coefficient) | z | p-value |
| --- | --- | --- | --- | --- | --- | --- |
| Mean RLTL Change | -5.021 (1.372) | 0.007 | 0.0004-0.097 | 151.572 | -3.659 | <0.001 |
| RLTL at the age of 1 year | -0.882 (0.479) | 0.414 | 0.162-1.059 | 2.415 | -1.840 | 0.066 |
| Average lifetime milk production | -7.798 *10<sup>-5</sup> (2.914 *10<sup>-5</sup>) | 1.000 | 1.000-1.000 | 1.000 | -2.676 | 0.007 |

## Supplementary File 2

### Detailed Materials and Methods

#### Animal population and data collection

At the Crichton Royal Farm, 200 milking cows plus their calves and replacement heifers are kept at any time. One half of the milking cows belong to a genetic line that has been selected for high milk protein and fat yield (S), while the other half is deliberately maintained on a UK average productivity level (C). Selection for these two genetic lines started in the 1970s. Animals of the C and S line do not significantly differ in their frame, weight or body condition score (p>0.05) as determined using t-tests for the animals in the present study. Each new-born calf is weighed and ear-marked and kept in an individual housing for the first few days of its life. Then C and S calves are transferred to outsides pens with a shelter, where groups of calves live together. All heifers are managed in the same way until their first calving when they are randomly allocated to a high forage (HF) or low forage (LF) diet. The LF diet is based on human food by-products and consists of a concentrate blend. The HF diet on the other hand is based on feed that is grown at the Crichton Royal Farm. While cows on a HF diet are turned out over the summer months for grazing and are only housed over the winter months, cows on a LF diet are housed continuously over the year without a grazing period. The food and water consumption of all calves and cows is monitored. All cows are milked three times daily and milk yield, milk composition and the milk somatic cell count as an indicator for (subclinical) mastitis are recorded. In the present study, these measurements were used to calculate an average milk production in kg per cow including all started lactations. The average of these measurements per cow across their lifetime was calculated and is referred to below as “average lifetime milk productivity” (Figure S3). Every day cows leave the milking parlour over a pressure plate which detects signs of lameness. Behaviour and health events are documented after visual detection by farm workers (Figure S4). At the end of the animal’s life its productive lifespan and a reason for culling is recorded. Productive lifetime is the time from birth to culling in days and is a proxy for the health span of the animal, because all animals that remain healthy enough to generate profit for the farmer remain in the herd. The most frequent reasons for culling were reproductive problems, mastitis, lameness and injuries caused by accidents (Figure S5). Along with a plethora of data that is recorded for each animal, routine blood sampling takes place initially shortly after birth and then annually in spring. Because of this sampling routine and because calves are being born all year round, age at sampling and sampling intervals vary for adult animals (Figure S3).

#### DNA extraction and qPCR

DNA from whole blood samples was extracted with the DNeasy Blood and Tissue spin column kit (QIAGEN). DNA samples had to have a minimum yield, purity and integrity to pass our internal quality control. Yield and purity were measured on a NanoDrop ND-1000 spectrophotometer (Thermo Scientific) and the DNA integrity was evaluated on integrity gels following Seeker et al. (2016) (1). In total, 1,328 samples of 308 animals with a minimal concentration of 20 ng/μl and ratios of 260/280 > 1.7 and 260/230 >1.8 that also had a DNA integrity score of 1 or 2 (1) were included on qPCR plates for RLTL measurements. We measured telomeric DNA in relation to the reference gene beta-2-microglobulin (B2M) that is constant in copy number (1) and has been used before in telomere studies on ruminant species such as Soay sheep, Roe deer and dairy cattle (1–5). Both reactions were performed on the same qPCR plate but in different wells (monoplex qPCR). An identical sample was included as a calibrator (or “golden sample”) twice on all 25 qPCR plates: one time in the middle of the plate and another time at its periphery. The measurements for the calibrator sample were used in the calculation of RLTL of individual samples to correct for part of the random measurement error that was associated with the qPCR plate. The two locations of the calibrator were used to test for a qPCR plate edge effect. All other samples were randomly allocated to qPCR plates and wells. Also, a negative control (DNAse and RNase free water) and a serial dilution of the calibrator DNA were added to each qPCR plate for visual qPCR quality control. A liquid handling robot (Freedom Evo by TECAN) was used to load samples, the calibrator, the negative control and the serial dilution in triplicates onto 384 well qPCR plates. For the amplification of telomeres tel 1b (5’-CGG TTT GTT TGG GTT TGG GTT TGG GTT TGG GTT TGG GTT-3’) and tel 2b (5’-GGC TTG CCT TAC CCT TAC CCT TAC CCT TAC CCT TAC CCT-3’) primers were used (6). B2M primers were obtained from (Primerdesign, accession code NM_001009284).

The following qPCR protocol was used on a LightCycler 480 (Roche): 15 min at 95 °C for enzyme activation followed by 50 cycles of 15 s at 95 °C (denaturation), 30 s at 58 °C (primer annealing) and 30 s at 72 °C (signal acquisition). The melting curve was acquired as follows: 1 min at 95 °C, followed by 30 s at 58 °C and a continuous increase of 0.11 °C/s to 95 °C with continuous signal acquisition.

The software LinReg PCR (7) was used for fluorescence baseline correction of raw RLTL measurements and for the calculation of reaction specific qPCR efficiencies for each plate (ETEL and EB2M for the telomere and the B2M reaction respectively). The following formula was used for RLTL calculation (8):

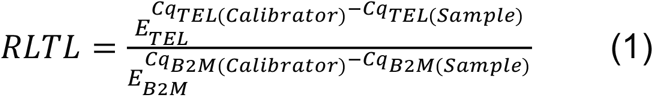

The Cq value describes the number of cycles of a qPCR that is required for an amplification curve to cross a set fluorescence threshold. The Cq values of the calibrator sample were Cq_TEL(Calibrator)_ and Cq_B2M(Calibrator)_ for the telomere and the B2M reaction respectively. Cq values of the individual samples were Cq_TEL(Sample)_ and Cq_B2M(Sample)_.

#### Statistical analysis

We have shown before that our RLTL data are significantly affected by qPCR plate and qPCR row (1, 9). To account for those known sources of measurement error, we used the residuals of a linear model that corrected all RLTL measurements for qPCR plate and row, by fitting plate and row as fixed factors in the model. These residual RLTL measures were used in all subsequent calculations and models of telomere dynamics. RLTL change was calculated as the difference between two subsequent adjusted RLTL measurements within individuals (RLTL change = RLTLt - RLTLt-1). To investigate which factors affect the direction and amount of RLTL change, a linear mixed model was fitted with RLTL change as response variable and animal identity as random effect. The following factors were included as fixed effects in the model to test their impact on RLTL change: genetic line, feed group and birth year of the animal, age at sampling (at time t), and the occurrence of a health event within two weeks before or after sampling (at time t). The time difference between consecutive samplings in days was fitted as a covariate. Non-significant fixed effects (p>0.05) were backwards eliminated from the model. Age at sampling was modelled as a covariate (age in years). Because we wanted to investigate if milk productivity was associated with RLTL change, the model of RLTL change was repeated for 918 change measurements of 253 animals that had milk productivity measurements available and average lifetime milk production was fitted as additional covariate.

To investigate the association between RLTL change and productive lifespan we first focussed on RLTL change within the first year of life by only considering two RLTL measurements per animal: The first was taken shortly after birth and the second at the approximate age of one year. The change between those measurements was tested as an explanatory variable in a cox proportional hazard model of productive lifespan. Then we investigated the association between lifetime telomere length dynamics and productive lifespan and calculated for each animal: 1) mean RLTL across all samples per animal 2) mean RLTL change and 3) mean absolute RLTL change. We tested these measures for lifetime telomere length dynamics fitted separately and together in cox proportional hazard models of productive lifespan that also included mean milk production as covariate. We have reported before that on average RLTL shortens in this population of dairy cattle within the first year of life while remaining on average stable thereafter (9, 10). Therefore, we were concerned that the initial RLTL attrition may contribute to more overall RLTL shortening in short-lived animals with less follow up samples. Therefore, we tested all models again while excluding all measurements that were taken shortly after birth and then also while excluding all animals with less than three RLTL measurements. Lastly, we tested if the previously reported effect of RLTL at the age of one year (10) remained statistically significant when tested in the same cox proportional hazard model as milk productivity and mean RLTL change.

All statistical analyses were performed in R studio with R 3.1.3. (11). Mixed-effects models were implemented using the ‘lme4’ library (version 1.1-18-1) (12) and figures were generated with the library ‘ggplot2’ (version 3.1.0) (13). For the Cox proportional hazard models the R package ‘survival’ (version 2.43-1) was used and for plotting survival curves we used ‘survminer’ (version 0.4.3).

